# The host factor ANP32A is required for influenza A virus vRNA and cRNA synthesis

**DOI:** 10.1101/2021.04.30.442228

**Authors:** Benjamin E. Nilsson-Payant, Benjamin R. tenOever, Aartjan J.W. te Velthuis

## Abstract

Influenza A viruses are negative-sense RNA viruses that rely on their own viral replication machinery to replicate and transcribe their segmented single-stranded RNA genome. The viral ribonucleoprotein complexes in which viral RNA is replicated consist of a nucleoprotein scaffold around which the RNA genome is bound, and a heterotrimeric RNA-dependent RNA polymerase that catalyzes viral replication. The RNA polymerase copies the viral RNA (vRNA) via a replicative intermediate, called the complementary RNA (cRNA), and subsequently uses this cRNA to make more vRNA copies. To ensure that new cRNA and vRNA molecules are associated with ribonucleoproteins in which they can be amplified, the active RNA polymerase recruits a second polymerase to encapsidate the cRNA or vRNA. Host factor ANP32A has been shown to be essential for viral replication and to facilitate the formation of a dimer between viral RNA polymerases and differences between mammalian and avian ANP32A proteins are sufficient to restrict viral replication. It has been proposed that ANP32A is only required for the synthesis of vRNA molecules from a cRNA, but not vice versa. However, this view does not match recent molecular evidence. Here we use minigenome assays, virus infections, and viral promoter mutations to demonstrate that ANP32A is essential for both vRNA and cRNA synthesis. Moreover, we show that ANP32 is not only needed for the actively replicating polymerase, but also for the polymerase that is encapsidating nascent viral RNA products. Overall, these results provide new insights into influenza A virus replication and host adaptation.

**IMPORTANCE:** Zoonotic avian influenza A viruses pose a constant threat to global health and they have the potential to cause highly pathogenic pandemic outbreaks. Species variations in host factor ANP32A play a key role in supporting the activity of avian influenza A virus RNA polymerases in mammalian hosts. Here we show that ANP32A acts at two stages in the influenza A virus replication cycle, supporting recent structural experiments and in line with its essential role. Understanding how ANP32A supports viral RNA polymerase activity and how it supports avian polymerase function in mammalian hosts is important for understanding influenza A virus replication and the development of antiviral strategies against influenza A viruses.

## INTRODUCTION

Influenza A viruses (IAV) are segmented negative-sense single-stranded RNA viruses that belong to the *Orthomyxoviridae.* IAV viruses infect a wide range of host species including humans, with wild aquatic birds being their natural reservoir (1). Humans are typically infected with seasonal, mammalian adapted IAV strains and only occasionally infected with avian adapted IAV strains. This is in part because zoonotic avian IAV strains are unable to efficiently transmit to and replicate in mammalian cells, unless multiple host species barriers are overcome (2). The two most important adaptations are i) a change in receptor binding specificity from α-2,3-linked sialic acids to α-2,6-linked sialic acids for effective viral entry into human cells (1), and ii) restored binding of the viral RNA-dependent RNA polymerase (RdRp) of avian IAV strains to host factor ANP32A for improved viral replication (3, 4).

The IAV RNA polymerase is composed of three subunits called polymerase basic 1 (PB1), PB2, and polymerase acidic (PA). The RNA polymerase replicates and transcribes the IAV genome segments in the context of viral ribonucleoprotein (vRNP) complexes (Fig 1A and 1B). In these vRNPs, the RNA polymerase binds to the ends of a single-stranded negative-sense viral RNA genome segment, while the viral nucleoprotein (NP) associates with the rest of the viral RNA (vRNA) segment in a double helical structure (5). During viral genome replication, vRNA segments are replicated by the viral RdRp through a complementary positive-sense RNA (cRNA) replicative intermediate, which requires at least one additional RNA polymerase to encapsidate the cRNA into a cRNP (5) (Fig 1A). The RNA polymerase in the cRNP next uses the cRNA as template to produce new vRNA genome segments, which is a process that requires at least one additional RdRp complex acting in a *trans*-activating capacity (5), and an RNA polymerase that encapsidates the vRNA into a new vRNP (6, 7) (Fig 1B).

**FIG 1.**
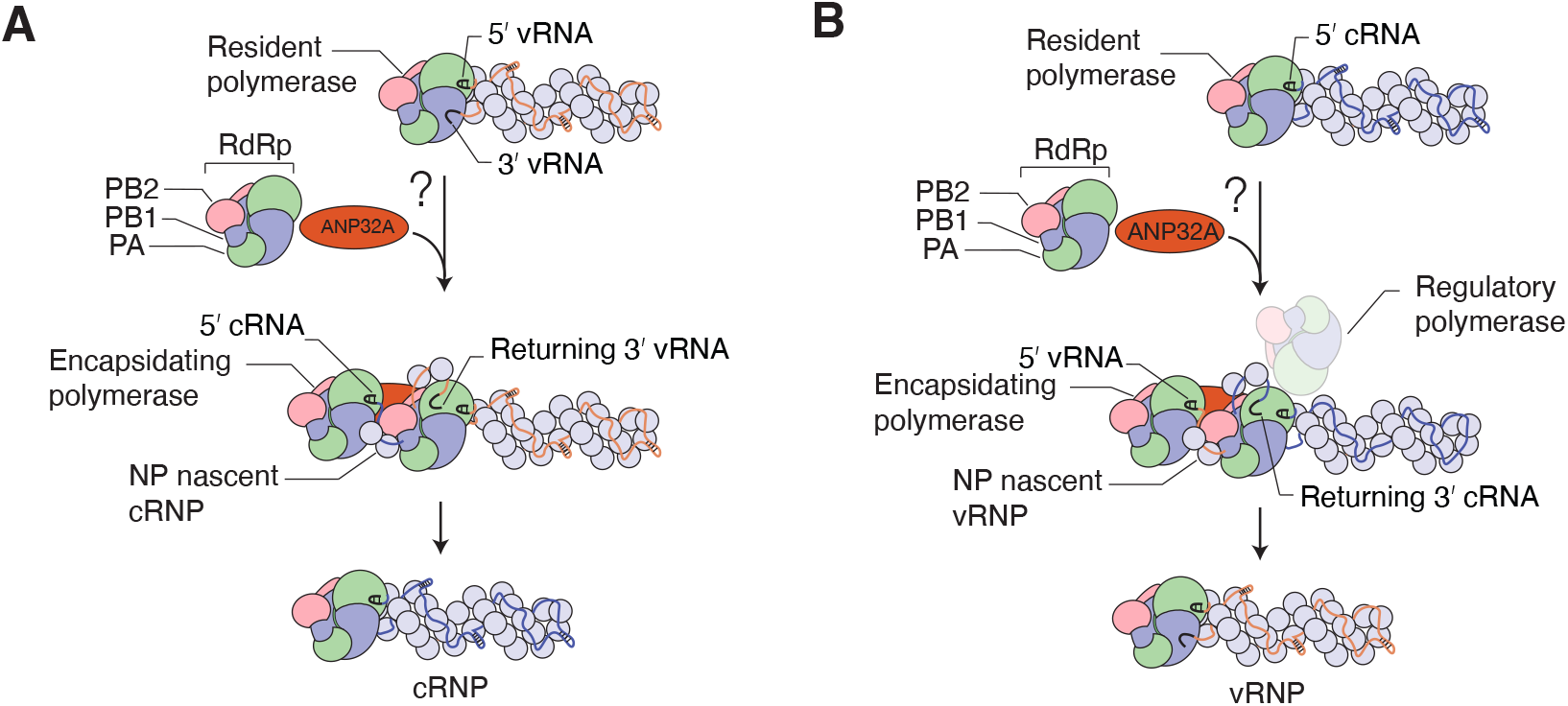
Influenza A virus genome replication. **(A)** Schematic of vRNA-to-cRNA replication by the resident RNA polymerase in vRNPs. Nascent cRNA is encapsidated by a newly synthesized RNA-free polymerase. **(B)** Schematic of cRNA-to-vRNA replication by the resident RNA polymerase aided by a second regulatory polymerase. Nascent vRNA is being encapsidated by a newly synthesized RNA-free polymerase.

One or more steps of avian-adapted IAV replication process are severely impaired in mammalian cells (2, 8), but a single glutamic acid-to-lysine mutation in the PB2 subunit (PB2-E627K) is sufficient to restore the activity of avian RNA polymerases in mammalian host cells (4). The molecular details underlying the impaired activity of avian RNA polymerases in mammalian cells are not fully understood, but several potential mechanisms have been proposed, including destabilized interactions of RNA polymerase and NP during vRNP assembly (9–12), reduced viral promoter RNA binding (13, 14), differential interactions with importin-α (15, 16), inhibitory or activating host factors (17, 18) and unstable cRNP structures of avian IAV in mammalian cells (19). However, expression of acidic (leucine-rich) nuclear phosphoprotein 32 family member A protein (ANP32A) from avian cells alone is sufficient to restore avian IAV RNA polymerase activity in mammalian cells (20), suggesting that the PB2-E627K mutation arises to compensate for an impaired interaction between mammalian ANP32A and the avian IAV RNA polymerase.

ANP32A consists of an N-terminal leucine-rich repeat (LRR) domain with four LRR motifs and a C-terminal low-complexity acidic region (LCAR) (21–23). Although the avian and mammalian ANP32A homologues have a high sequence identity, birds contain an additional exon duplication (24, 25). Alternative splicing of the avian ANP32A gene leads to three ANP32A proteins: one containing an additional SUMO interaction motif (SIM)-like and LCAR sequence encoded by an additional exon; one containing an additional LCAR sequencing through partial splicing of the additional exon; and one lacking the extra exon and being similar to the mammalian ANP32A protein (25–27). Restoring avian IAV RdRp activity in mammalian host cells by avian ANP32A overexpression was shown to be dependent on the presence of the avian-specific exon duplication (20). ANP32A’s closely related family member ANP32B, which is required for mammalian cell-adapted RdRp function (28, 29), does not appear to play a role in restoring replication of avian-adapted IAV (30).

Interaction studies of ANP32A and the viral RNA polymerase have suggested that ANP32A only interacts with the complete heterotrimeric RNA polymerase (27, 28, 31, 32). ANP32A proteins containing the avian signature exon duplication were also shown to have a much stronger affinity for the viral RNA polymerase (26, 27, 33). Presence of the avian glutamic acid residue 627 (627E) on the PB2 subunit significantly weakens the interaction with ANP32A, necessitating the additional ANP32A exon or PB2 mutation E627K to stabilize the interaction (34). The determinant of the differential interaction was mapped to the C-terminal LCAR domain of ANP32A, which directly interacts with the flexible PB2 627-domain on the viral RdRp (27, 33).

Interestingly, the interaction between ANP32A and the viral RNA polymerase is significantly strengthened in the presence of viral RNA (26, 33), suggesting that ANP32A binds the RNA polymerase in the context of an RNP. Recent cryo-electron microcopy structures of an influenza C virus RNA polymerase dimer revealed that ANP32A formed a bridge between an RNA polymerase bound to RNA and an apo form of the RNA polymerase (7), suggesting that ANP32A mediates the assembly of the viral replicase complex. In line with this idea, ANP32A appears to be required for viral RNA genome replication, but not primary transcription, in both avian and mammalian hosts (20, 28). Moreover, using recombinant purified protein, human ANP32A has been shown to be an enhancer of vRNA synthesis by human-adapted IAV RNA polymerases (28). Based on the latter observation, it has been proposed that ANP32A is only required for the synthesis of vRNA molecules from a cRNA template, but not vice versa. However, both cRNA and vRNA molecules are encapsidated and it is unlikely that IAV evolved separate encapsidation complexes for vRNA and cRNA nascent strands.

To investigate at which stage in IAV replication ANP32A is critical, we used a combination of cell-based minigenome assays, single-step promoter mutants, and recombinant IAV strains to systematically characterize the role of ANP32A in IAV genome replication. We find that an IAV RNA polymerase with a 627E mutation is unable to efficiently synthesize both vRNA and cRNA, and that expression of avian ANP32A is sufficient for the 627E RNA polymerase to restore both vRNA and cRNA synthesis. Overall, this study provides new insights into the role of ANP32A in mediating IAV replicase function. Together these insights lead to a better understanding of IAV genome replication and host adaptation.

## RESULTS

### Avian ANP32A stimulates infection and replication of IAV containing PB2 627E

Avian IAV RNA polymerase activity is significantly restricted in mammalian host cells (14, 17, 35) and it has been demonstrated that the expression of avian host protein ANP32A, but not ANP32B, is sufficient to restore avian IAV RNA polymerase activity in mammalian cells (20, 30). However, the exact step at which ANP32A is involved in RNA polymerase activity remains unclear. We set out to systematically analyze the role of ANP32A using PB2 627E as tool and exchanged the lysine of the mammalian-adapted influenza strain A/WSN/33 (H1N1) (abbreviated as WSN) with the avian-specific glutamic acid (WSN-K627E) using reverse genetics. As has been reported previously (17), the avian-like WSN-K627E virus was significantly restricted in both viral replication and transcription in human cells compared to WSN (Fig 2A). However, in avian cells, no significant difference in viral replication and transcription was observed between both viruses. To confirm that infection of human cells with the avian-like WSN-K627E virus would be restored by avian ANP32A, we transiently expressed chicken ANP32A (chANP32A) in HEK-293T cells and infected the cells with either wild-type WSN or avian-like WSN-K627E virus. While WSN-K627E was unable to grow in human cells in the absence of avian ANP32A, expression of chANP32A was sufficient to support WSN-K627E growth similar to the wild-type WSN virus (Fig 2B). Moreover, analysis of the accumulation of viral RNA in human cells showed that transient expression of chANP32A, but not human ANP32A (huANP32A), was able to fully restore restricted viral RNA synthesis by WSN-K627E, while exogenous expression of either chANP32A or huANP32A had no discernible effect on the replication of the wild-type WSN (Fig 2C).

**FIG 2.**
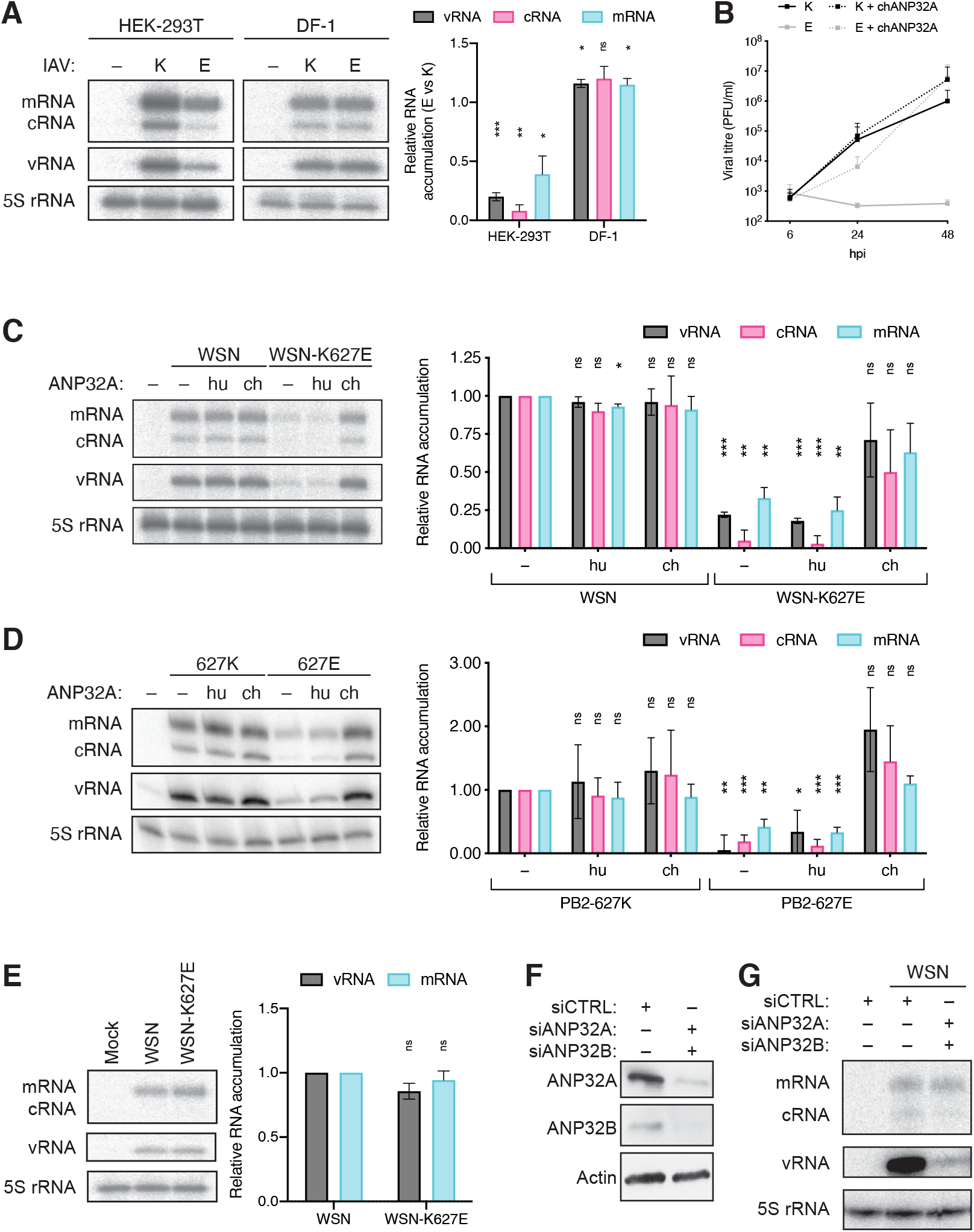
ANP32A is required for IAV genome replication. **(A)** Human HEK-293T cells or chicken DF-1 cells were infected with influenza A/WSN/33 (H1N1) virus carrying either a lysine (K) or glutamic acid (E) on residue 627 of the PB2 subunit for 6 h at an MOI of 1. The accumulation of viral RNA was analyzed by primer extension and 6% PAGE. The graph shows the relative mean intensity of viral RNA normalized to 5S rRNA from three independent biological replicates. **(B)** HEK-293T cells were transiently transfected with plasmids expressing GFP or chANP32A. 24 h post-transfection, cells were infected with mammalian-(K) or avian-(E) adapted influenza A/WSN/33 (H1N) at an MOI of 0.01. Infectious viral titers were determined by plaque assay at the indicated timepoints post-infection from three independent biological replicates. **(C)** HEK-293T cells were transfected with plasmids expressing huANP32A or chANP32A or GFP as a negative control. 24h post-transfection, cells were infected with wild-type influenza A/WSN/33 (H1N1) virus (WSN) or virus with a PB2-K627E mutation (WSN-K627E) at an MOI of 1 for 6 h. The accumulation of viral RNA was analyzed by primer extension and 6% PAGE. The graph shows the relative mean intensity of viral RNA normalized to 5S rRNA from three independent biological replicates. **(D)** HEK-293T cells were transfected with plasmids expressing NP, PA, PB1, PB2 (627K or 627E), segment 6 vRNA and huANP32A, chANP32A or GFP. 48h post-transfection, the accumulation of viral RNA was analyzed by primer extension and 6% PAGE. The graph shows the relative mean intensity of viral RNA normalized to 5S rRNA from three independent biological replicates. **(E)** HEK-293T cells were infected with wild-type influenza A/WSN/33 (H1N1) virus (WSN) or virus with a PB2-K627E mutation (WSN-K627E) at an MOI of 10 for 6 h in the presence of cycloheximide. The accumulation of viral RNA was analyzed by primer extension and 6% PAGE. The graph shows the relative mean intensity of viral RNA normalized to 5S rRNA from three independent biological replicates. **(F-G)** A549 cells were transfected with siRNA targeting human ANP32A and ANP32B. 48 h post-transfection cells were either **(F)** lysed for Western blot analysis for ANP32A and ANP32B expression or **(G)** infected with influenza A/WSN/33 (H1N1) virus (WSN) at an MOI of 1 for 12 h. The accumulation of viral RNA was analyzed by primer extension and 6% PAGE. Error bars represent the standard deviation of the means and asterisks represent a significant difference from the control group (two-tailed one-sample t test) as follows: ns, *P* > 0.05; *, *P* < 0.05; **, *P* < 0.01; ***, *P* < 0.001; ****, *P* < 0.0001.

### Avian ANP32A stimulates replication, but not transcription by 627E-containing RNA polymerases

In order to confirm that the above observations are linked to the activity of the viral RNA polymerase and no other viral factors, we transiently reconstituted the minimal viral components necessary for genome transcription and replication in HEK-293T cells and analyzed the accumulation of the three viral RNA species cRNA, vRNA and mRNA. In agreement with the above findings, we found that vRNPs containing the PB2-627E subunit were significantly restricted in their activity, showing reduced production of all viral RNA species, and that viral RNA accumulation was restored in the presence of chANP32A (Fig 2D).

Accumulation of viral mRNA products is dependent on viral replication, which can confound analysis of the effect of ANP32A. As cellular protein translation is blocked by cycloheximide and protein translation is necessary for viral genome replication, but not transcription, cycloheximide can be used to obtain insight into primary transcription of incoming infecting vRNPs. To confirm that primary transcription by the 627E IAV vRNPs was independent of ANP32A in mammalian cells, we infected human HEK-293T cells at a high MOI with the WSN or WSN-K627E virus in the presence of cycloheximide. In agreement with previous reports, we found that there was no difference in mRNA synthesis, suggesting that the previously observed reduced mRNA levels were a consequence of impaired genome replication and consequently reduced vRNA template levels (Fig 2E).

The above result indicates that avian ANP32A is only required for the genome replication of avian-like IAVs and not for their transcription. However, the findings do not prove that mammalian ANP32A plays no role in the transcription of mammalian-adapted IAV RNA polymerases. Therefore, we knocked down both human ANP32A and ANP32B, which have been shown to be redundant in mammalian cells (29), in human alveolar epithelial A549 cells, and infected these cells with WSN at a high MOI. We found that early in the infection cycle, no difference in primary transcription was observed in control and knockdown cells, while a strong reduction in viral genome replication could be seen (Fig 2F and 2G). Overall, our findings suggest that ANP32A is only involved in IAV replication and they confirm previous reports that an IAV RNA polymerase with a 627E residue requires the presence of the avian ANP32A host factor to perform efficient genome replication, but not transcription.

### The N-terminal third of ANP32A LCAR domain is required for efficient IAV replication

An exon duplication in the avian ANP32A gene was linked to the stimulation of avian IAV RNA polymerase (20, 24, 25). More recently, the central domain of ANP32A immediately upstream of the avian exon duplication was linked to avian IAV RNA polymerase stimulation as well (30). In order to gain a better understanding of the role of the ANP32A domains in IAV replication, internal and C-terminal deletions were introduced into a chANP32A expression construct (Fig 3A and 3B). The ability of these truncated chANP32A proteins to enhance viral replication by an avian-like RNA polymerase was determined using minigenome assays (Fig 3C). Deletion of the exon duplication in the chANP32A protein led to a loss in 627E RNA polymerase activity relative to transient expression of the full-length chANP32A. However, in the presence of the exon duplication a C-terminal or an internal truncation of 31 amino acids in the LCAR domain was compatible with 627E RNA polymerase activity, while longer deletions of the LCAR domain led to a loss in 627E RNA polymerase activity. Together, these results suggest that the LCAR domain of ANP32A plays a pivotal role in supporting IAV RNA polymerase activity.

**FIG 3.**
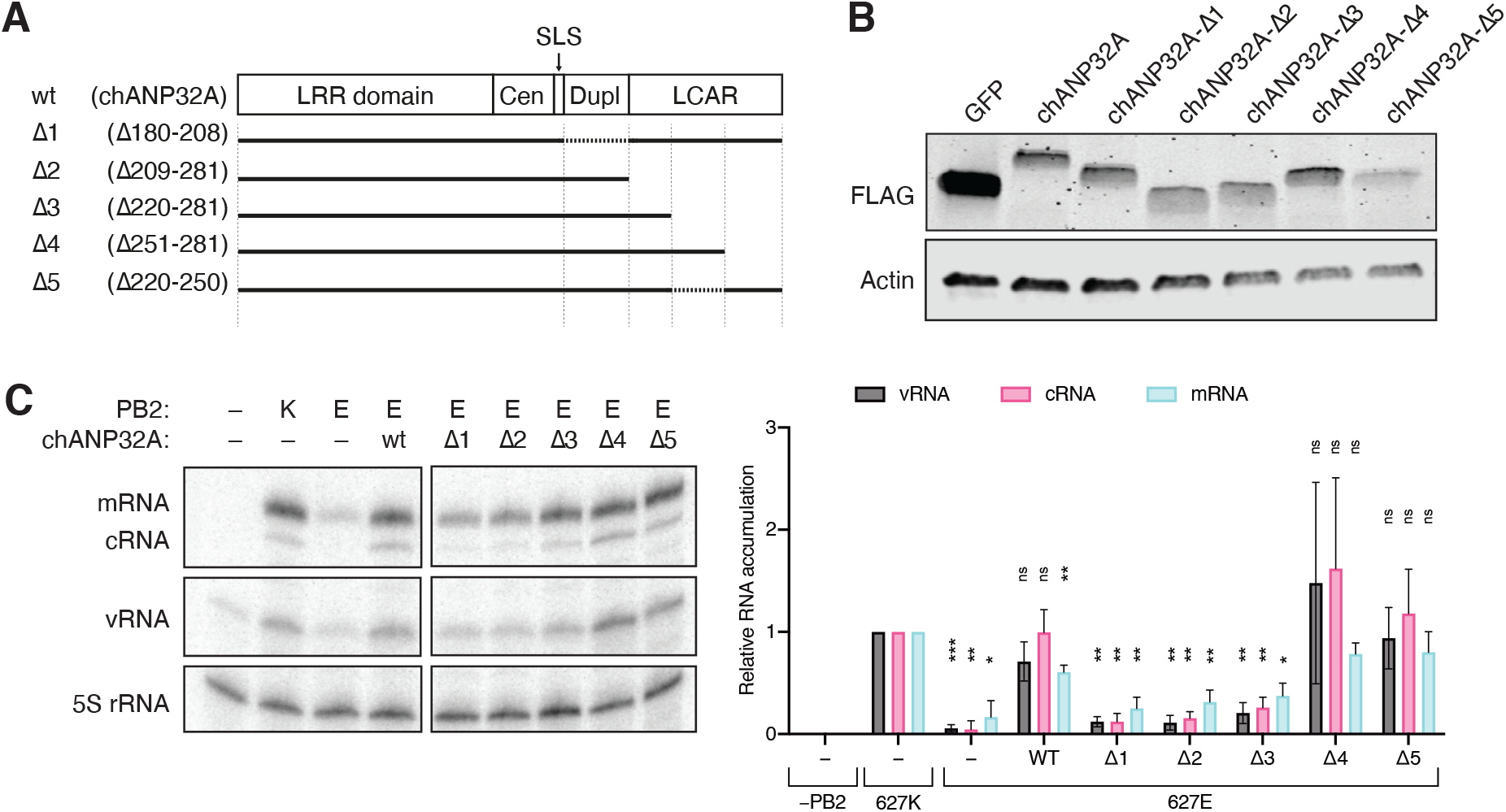
The chANP32A LCAR domain is essential for avian RdRp stimulation. **(A)** Schematic of the wild-type chANP32A protein and the internal and terminal truncations introduced in this study. **(B)** HEK-293T cells were transfected with plasmids expressing NP, PA, PB1, PB2 or PB2-K627E, segment 6 vRNA and either wild-type or truncated chANP32A. 48h post-transfection, the accumulation of viral RNA was analyzed by primer extension and 6% PAGE. The graph shows the relative mean intensity of viral RNA normalized to 5S rRNA from three independent biological replicates. Error bars represent the standard deviation of the means and asterisks represent a significant difference from the control group (two-tailed one-sample t test) as follows: ns, *P* > 0.05; *, *P* < 0.05; **, *P* < 0.01; ***, *P* < 0.001; ****, *P* < 0.0001.

### Promoter binding is not restricted in avian RNPs

Some reports have suggested that an avian RdRp forms inherently weaker interactions with the vRNA template than a mammalian adapted IAV RdRp (13, 14). The 627E residue is located far away from the promoter binding site and our previous work has shown that the 627-domain does not affect the basic functions of the RNA polymerase (36). In an effort to exclude that a single point mutation in PB2-627 leads to a significant difference in vRNA promoter binding and possible vRNP assembly, the IAV RNA polymerase, either carrying a PB2-627K (K) or PB2-627E (E) subunit, a catalytically inactive PB1 subunit (D445A/D446A) and a TAP-tagged PA subunit, was transiently expressed in human HEK-293T cells together with a 76 nucleotide-long vRNA based on segment 5. Following immunoprecipitation of the RNA polymerase, bound RNA was extracted and the amount of co-immunoprecipitated vRNA was determined (Fig 4A). We observed that vRNA binding of a PB2-627E carrying RNA polymerase was not impaired compared to a PB2-627K RNA polymerase. In addition, expression of chANP32A did not have any impact on vRNA binding.

**FIG 4.**
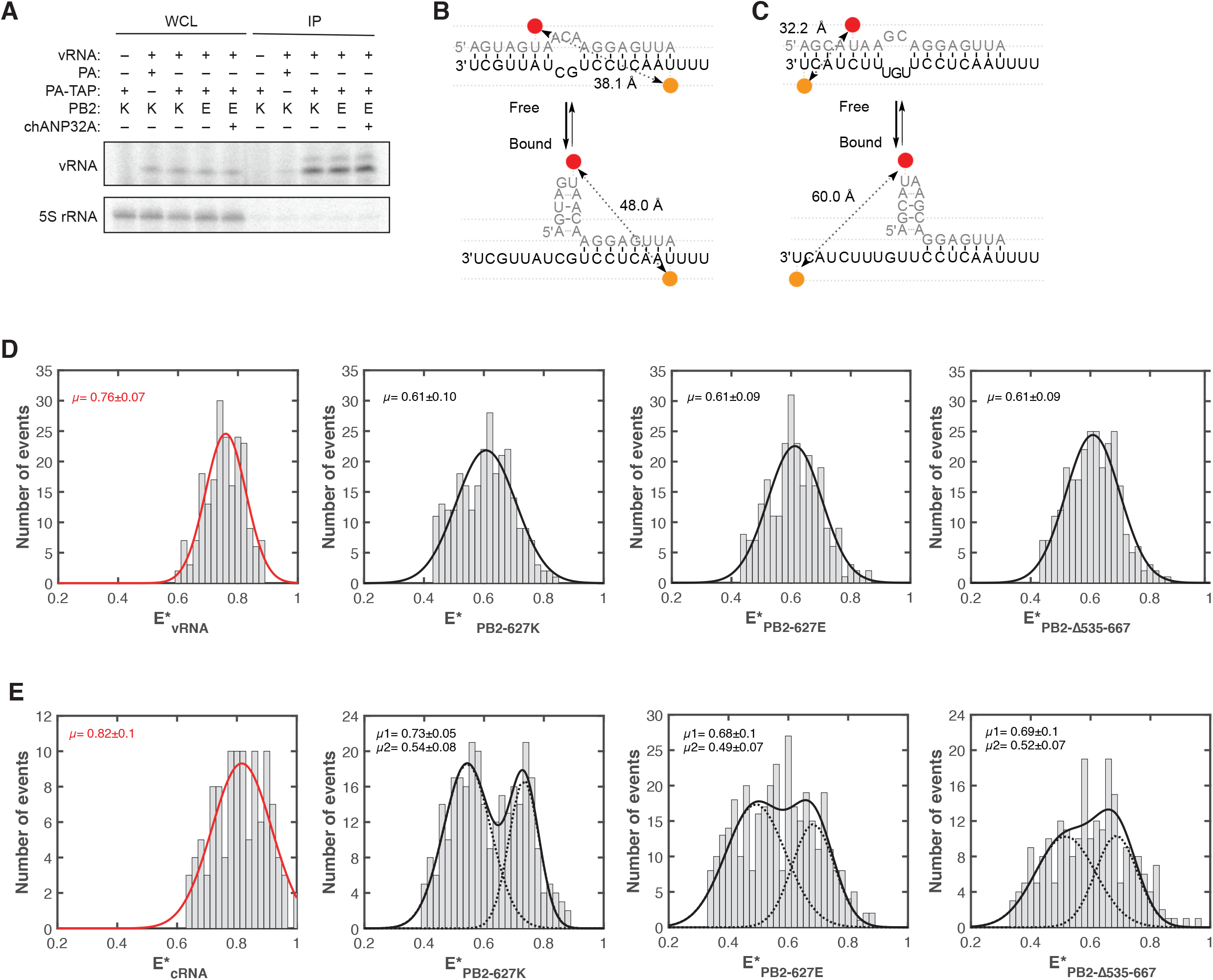
The PB2 627-domain does not affect viral RNA binding. **(A)** HEK-293T cells were transfected with plasmids expressing PA, PB2 (627K or 627E), catalytically inactive PB1, a 47 nucleotide-long internally truncated segment 6 and where indicated chANP32A. 48 h post-transfection, cells were lysed and the viral RdRp was immunoprecipitated via a TAP-tag on PA. Co-precipitated vRNA was analyzed by primer extension and 12% PAGE. **(B)** Schematic of the influenza A virus vRNA promoter before and after binding with the viral RdRp. The distance between the Atto647N dye (red) and the Cy3 dye (orange) is indicated. **(C)** Schematic of the influenza A virus cRNA promoter before and after binding with the viral RdRp. The distance between the Atto647N dye (red) and the Cy3 dye (orange) is indicated. **(D)** vRNA promoter binding by the influenza A virus RdRp as analyzed by smFRET. Recombinant viral RdRp, either with a PB2 subunit with a lysing (627K) or glutamic acid (627E) at position 627 or with a subunit lacking the entire 627-domain (Δ535-667) was used. The FRET populations were fit with a single Gaussian curve. The RNA only (red) is shown as comparison. The means of the bound (black dotted line) and unbound (red dotted line) signal is indicated in each graph. **(E)** cRNA promoter binding by the influenza A virus RdRp as analyzed by smFRET. Recombinant viral RdRp, either with a PB2 subunit with a lysing (627K) or glutamic acid (627E) at position 627 or with a subunit lacking the entire 627-domain (Δ535-667) was used. The FRET populations were fit with a double Gaussian curve representing the two conformations that the cRNA promoter adopts in a pre-initiation state. The unbound RNA only (red line) is shown as comparison.

To get a more quantitative insight into vRNA and cRNA promoter binding activity of the mammalian or avian-adapted IAV RNA polymerase, we performed single-molecule Förster resonance energy transfer (sm-FRET) assays using recombinant catalytically inactive but RNA-binding competent RNA polymerases and short fluorescently labelled viral RNA promoters. In an unbound state, both vRNA and cRNA promoters form largely double-stranded RNA molecules (37). However, if bound by the IAV RNA polymerase, the 5′ and 3′ promoter ends are separated in space, which increases the distance between donor and acceptor dyes (Fig 4B and 4C). We observed that the vRNA-bound FRET populations were similar, irrespective of whether the RNA polymerase added to the vRNA carried a lysine or glutamic acid at position 627 or in fact lacked the entire 627-domain (Fig 4D). Moreover, similar FRET populations, with a clear shift from the unbound state, were observed for the cRNA promoter after addition of the different RNA polymerases (Fig 4E). Together, these data demonstrate that the 627-domain of the PB2 subunit does not play a role in vRNA or cRNA promoter binding and that the nature of position PB2-627 has no impact on RNA polymerase-RNA binding.

### ANP32A function in RNA polymerase activity is NP-independent

The interaction between PB2 and NP has been implicated several times as the cause for avian RNA polymerase restriction in mammalian cells (9–12). Previously, it was reported that avian RNA polymerase activity was less diminished or even not diminished on short viral RNA templates that do not require NP for viral replication (14). In order to determine whether ANP32A’s function is NP or vRNA template length dependent, we therefore transiently expressed the IAV RNA polymerase in the presence or absence of NP and chANP32A together with a short segment 5-based 76 nucleotide-long vRNA and analysed the accumulation of viral RNA in this NP-independent minigenome assay (Fig 5A). We find that even in this NP-independent replication assay, the PB2-K627E mutation led to a significant loss in RNA polymerase activity, which was restored in the presence of chANP32A. The absence or presence of NP did not affect these results, suggesting that NP does not play any major role in ANP32A function. Next, we tested whether using shorter vRNA templates affects ANP32A function. Using both a 47 nucleotide-long (Fig 5B) and a 30 nucleotide-long (Fig 5C) segment 6-based vRNA template we confirmed our observation that avian RNA polymerase restriction and the role of ANP32A in IAV RNA replication is independent of both template length and NP.

**FIG 5.**
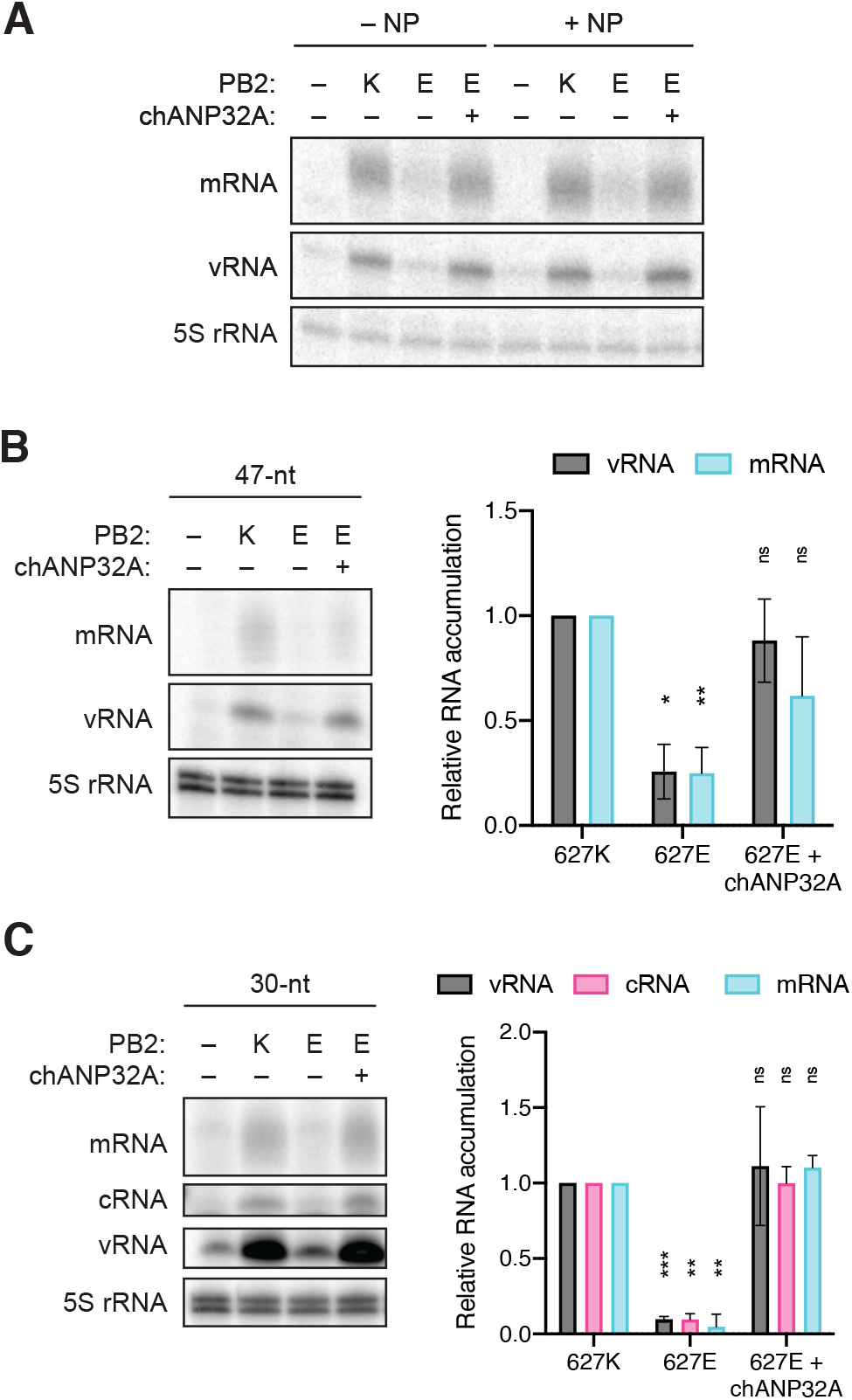
Avian RdRp restriction in mammalian cells is independent of NP. **(A)** HEK-293T cells were transfected with plasmids expressing a 76 nucleotide-long internally truncated segment 5 vRNA, PA, PB1, PB2 and where indicated NP and chANP32A. 48 h post-transfection, the accumulation of viral RNA was analyzed by primer extension and 12% PAGE. **(B)** HEK-293T cells were transfected with plasmids expressing a 47 nucleotide-long internally truncated segment 6 vRNA, PA, PB1, PB2 and where indicated chANP32A. 48 h post-transfection, the accumulation of viral RNA was analyzed by primer extension and 12% PAGE. The graph shows the relative mean intensity of viral RNA normalized to 5S rRNA from three independent biological replicates. **(C)** HEK-293T cells were transfected with plasmids expressing a 30 nucleotide-long internally truncated segment 6 vRNA, PA, PB1, PB2 and where indicated chANP32A. 48 h post-transfection, the accumulation of viral RNA was analyzed by primer extension and 12% PAGE. The graph shows the relative mean intensity of viral RNA normalized to 5S rRNA from three independent biological replicates. Error bars represent the standard deviation of the means and asterisks represent a significant difference from the control group (two-tailed one-sample t test) as follows: ns, *P* > 0.05; *, *P* < 0.05; **, *P* < 0.01; ***, *P* < 0.001; ****, *P* < 0.0001.

### ANP32A is required for both cRNA and vRNA synthesis in mini genome assays

Several studies have suggested that a pair of mutations in the 3′ vRNA promoter (G3A and C8U, Fig 6A) can improve avian RNA polymerase activity in mammalian cells (13, 14). We wondered if we could use the RNA polymerase promoter to uncouple vRNA synthesis and cRNA synthesis, and thus study at what step of viral replication ANP32A is essential. This can only be addressed using minigenome assays, since both cRNA and vRNA synthesis must occur during a viral infection. As a first step towards this aim, we performed minigenome assays using either a construct expressing wild-type segment 6 vRNA or a segment 6 vRNA containing these promoter mutations (3A8U), and found that viral replication was greatly enhanced by these promoter mutations compared to the wild-type vRNA template (Fig 6B). However, even at elevated levels, replication was still significantly, albeit less markedly, restricted by the 627E RNA polymerase. Additional expression of chANP32A fully restored the activity of the 627E RNA polymerase, in line with our observations above.

**FIG 6.**
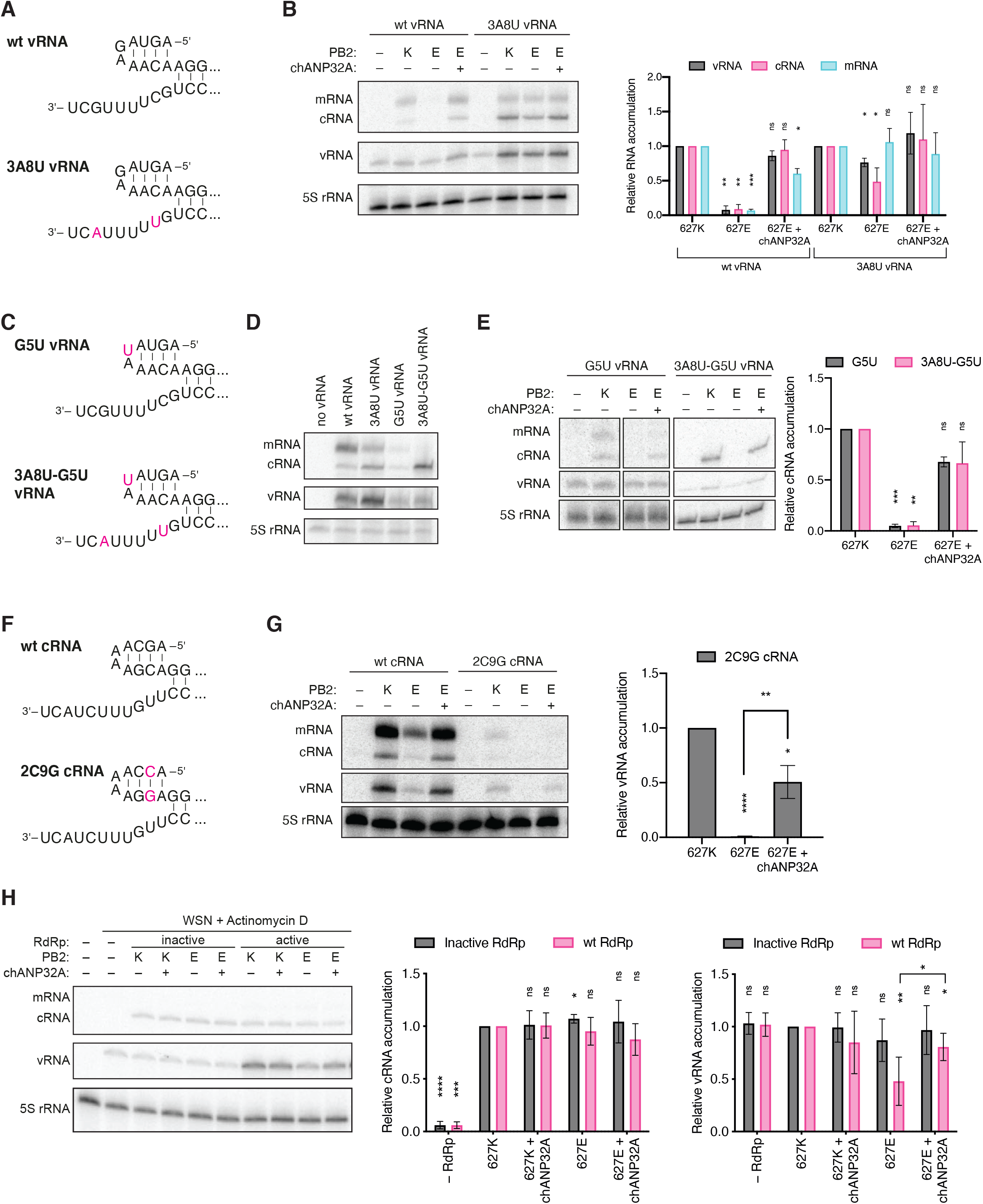
ANP32A is required for both vRNA and cRNA synthesis. **(A)** Schematic of the wild-type vRNA promoter or a vRNA promoter carrying a G3A and C8U mutation in the 3′ vRNA promoter (red). **(B)** HEK-293T cells were transfected with plasmids expressing NP, PA, PB1, PB2 (627K or 627E), segment 6 vRNA and chANP32A where indicated. 48h post-transfection, the accumulation of viral RNA was analyzed by primer extension and 6% PAGE. The graph shows the relative mean intensity of viral RNA produced from the mutant 3A8U vRNA template normalized to 5S rRNA from three independent biological replicates. **(C)** Schematic of the vRNA promoter carrying a G5U mutation in the 5′ vRNA promoter with and without the 3A8U 3′ vRNA promoter mutation. **(D)** HEK-293T cells were transfected with plasmids expressing NP, PA, PB1, PB2 (627K or 627E), segment 6 vRNA and chANP32A where indicated. 48h post-transfection, the accumulation of viral RNA was analyzed by primer extension and 6% PAGE. The graph shows the relative mean intensity of cRNA normalized to 5S rRNA from three independent biological replicates. **(E)** Schematic of the wild-type cRNA promoter or a cRNA promoter carrying a G2C and C9G mutation in the 5′ cRNA promoter. **(F)** HEK-293T cells were transfected with plasmids expressing NP, PA, PB1, PB2 (627K or 627E), segment 6 cRNA and chANP32A where indicated. 48h post-transfection, the accumulation of viral RNA was analyzed by primer extension and 6% PAGE. The graph shows the relative mean intensity of vRNA produced from the mutant 2C9G cRNA template normalized to 5S rRNA from three independent biological replicates. **(G)** HEK-293T cells were transfected with NP, PA, PB1 or catalytically inactive PB1a, PB2 (627K or 627E) and chANP32A where indicated. 24 h post-transfection, cells were infected with influenza A/WSN/33 (H1N1) virus at an MOI of 5 for 6 h in the presence of Actinomycin D. The accumulation of viral RNA was analyzed by primer extension and 6% PAGE. The graph shows the relative mean intensity of viral RNA normalized to 5S rRNA from four independent biological replicates. Error bars represent the standard deviation of the means and asterisks represent a significant difference from the control group (two-tailed one-sample t test) or between two non-control groups (two-tailed two-sample t test) as follows: ns, *P* > 0.05; *, *P* < 0.05; **, *P* < 0.01; ***, *P* < 0.001; ****, *P* < 0.0001.

Next, we introduced an additional mutation in the conserved 5′ vRNA stem-loop (G5U, Fig 6C) to address whether the 3A8U mutation affects cRNA synthesis. The G5U mutation in the 5′ vRNA promoter prevents internal initiation on the resulting cRNA product and therefore only allows primary mRNA as well as primary cRNA synthesis but block subsequent vRNA synthesis and secondary transcription. This becomes especially apparent when combining the G5U and 3A8U mutations (Fig 6D). Interestingly, minigenome assays with vRNA templates carrying the G5U 5′ vRNA promoter mutation clearly demonstrated that the 627E RNA polymerase mutation prevented efficient primary cRNA synthesis. Activity of the 627E RNA polymerase was restored by the additional expression of chANP32A (Fig 6E). These observations were evident when using both the wild-type or the 3A8U 3′ vRNA promoter, lending further proof to our finding that the 3A8U promoter mutation alone is not sufficient to overcome the restriction of the avian-like 627E RNA polymerase.

In order to determine whether only cRNA synthesis or also vRNA synthesis is restricted in a 627E RNA polymerase and whether ANP32A is required for efficient vRNA synthesis, we introduced a pair of mutations (2C9G) in the 5′ cRNA promoter that would prevent 3’ terminal initiation on the resulting vRNA product, thereby abolishing further cRNA synthesis, but not transcription (Fig 6F). Indeed, in a minigenome assay no vRNA synthesis could be observed in the presence of the 2C9G cRNA template and the 627E RNA polymerase (Fig 6G). Additional expression of chANP32A again was able to significantly increase RNA polymerase activity.

### ANP32A is required for both cRNA and vRNA synthesis during infection

In order to test whether these results could be recapitulated in the context of an infection and whether avian RNA polymerase restriction occurs at the level of the actively replicating RNA polymerase or at the level of a secondary RdRp necessary to form nascent RNPs, we assessed the ability of an incoming vRNP to synthesize cRNA, and of the resulting cRNP to perform further genome replication. Specifically, we pre-expressed either catalytically inactive (PB1a) or active (wt) RNA polymerase, NP and chANP32A as indicated, before infecting cells with WSN in the presence of the cellular transcription inhibitor Actinomycin D (Fig 6H). Therefore, viral replication is entirely reliant on the presence of pre-expressed viral components. In the absence of IAV RNA polymerase, no cRNA stabilization could be observed. Both PB2-627K or PB2-627E RNA polymerases were able to stabilize nascent cRNA equally well when active replication by pre-expressed RNA polymerases was prevented by the D445A/D446A mutation in the PB1 subunit. However, when further replication was enabled by pre-expression of a catalytically active RNA polymerase, reduced levels of vRNA synthesis were observed by the now actively replicating PB2-627E RNA polymerase, which was overcome by the addition of chANP32A.

Overall, these results suggest that IAV RNA polymerases containing a PB2-K627E mutation are impaired at the level of the actively replicating RNA polymerase and not the encapsidating RNA polymerase that is required for nascent RNA binding. Furthermore, it also demonstrates that avian ANP32A is only required for the actively replicating and not the encapsidating 627E RNA polymerase.

## DISCUSSION

The RNA polymerase activity of avian IAV is significantly impaired in mammalian host cells and it has been proposed that host factor ANP32A may restrict one step of the viral replication process. Here, we report that ANP32A is required for both efficient vRNA as well as cRNA synthesis, but not viral transcription. This is in agreement with previous studies (20, 28). In addition, we found that the N-terminal part of the LCAR of ANP32A is critical for successful IAV RNA polymerase stimulation, in line with previous reports (27, 38). Moreover, we show that the reduced activity of an IAV RNA polymerase containing PB2-627E in mammalian cells is independent of vRNA or cRNA promoter binding, in contrast with previous reports that implicated weaker promoter binding capabilities of avian-adapted RNA polymerases (13, 14). These findings clearly confirm that ANP32A plays a more fundamental role in IAV genome. In addition, while a previous study found that avian RdRp restriction was diminished or altogether abolished when using shorter vRNA templates, we found that an inability to replicate vRNA templates remained, even when using templates of 30 nucleotides in length (14).

Our finding that avian-adapted RNA polymerases are able to stabilize nascent cRNA products equally well in human host cells as mammalian-adapted RNA polymerases is in agreement with previous studies (19, 36). However, here we additionally demonstrate that avian-adapted cRNPs are unable to produce vRNA efficiently, suggesting that vRNA synthesis is impaired in the absence of the species-specific ANP32A host protein. Furthermore, we also illustrate that not only vRNA synthesis is restricted, but that in the context of minigenome assays, where only single-round replication cycles are permitted, both vRNA and cRNA synthesis are inhibited by the PB2-K627E mutation and subsequently restored by the expression of chANP32A. These data are consistent with recent structural findings where ANP32A is found to mediate the assembly of the IAV replicase complex consisting of an actively replicating vRNP bound by ANP32A which recruits an encapsidating RdRp (7).

In summary, these findings confirm that IAVs hijack ANP32A to facilitate viral genome replication by mediating assembly of the replicase complex. It is tempting to speculate that IAVs evolved this dependency in avian species. Because the mammalian ANP32A is shorter than its avian homologue, it cannot efficiently support assembly of the IAV replicase complex at both stages of viral replication. Zoonotic IAV strains must therefore acquire adaptive mutations to restore the ANP32A-RNA polymerase interaction. These findings strengthen previous hypotheses about IAV host range restriction and shed further light on the intricate details of IAV genome replication.

## MATERIALS AND METHODS

### Cell Culture

Human alveolar basal epithelial carcinoma cells (A549, ATCC, CCL-185), human embryonic kidney cells (HEK-293T, ATCC, CRL-3216), chicken embryonic fibroblasts (DF-1, ATCC, CRL-12203) and Madin-Darby Canine kidney epithelial cells (MDCK, ATCC, CCL-34) were commercially obtained. All cells were maintained at 37°C and 5% CO_2_ in Dulbecco’s Modified Eagle Medium (DMEM) supplemented with 10% Fetal Bovine Serum (FBS) and penicillin and streptomycin.

### Plasmids

Plasmids pcDNA-NP, pcDNA-PA, pcDNA-PB1, pcDNA-PB2, pcDNA-3a (45), pcDNA-PB1a (46), pcDNA-PB2-K627E (10), pcDNA-PB2Δ535-667 (36), pcDNA-PA-TAP (47), pPOLI-NA (48), pPOLI-cNA (49), pPOLI-NA47 (14), pPOLI-NP76 (50), as well as the pHW2000 constructs expressing the eight influenza A/WSN/33 (H1N1) virus segments (51) have been described previously.

Plasmids pCMV-3xFLAG-GFP, pCMV-3xFLAG-huANP32A and pCMV-3xFLAG-chANP32A were generated by InFusion cloning of coding sequences for GFP, huANP32A and chANP32A (synthesized by Integrated DNA Technologies) into an empty pCMV-3Tag-1A vector (Agilent Technologies).

Plasmids pHW2000-PB2-K627E, pPOLI-NA-3A8U, pPOLI-NA-G5U, pPOLI-NA-3A8U/G5U and pPOLI-cNA-2C9G were generated by site-directed PCR mutagenesis using the primers detailed in Table 1. Plasmids pPOLI-NA30, pCMV-3xFLAG-chANP32AΔ180-208, pCMV-3xFLAG-chANP32AΔ209-281, pCMV-3xFLAG-chANP32AΔ220-281, pCMV-3xFLAG-chANP32AΔ251-281 and pCMV-3xFLAG-chANP32AΔ220-250 were generated by deletion PCR mutagenesis using primers detailed in Table 1.

**Table 1.**
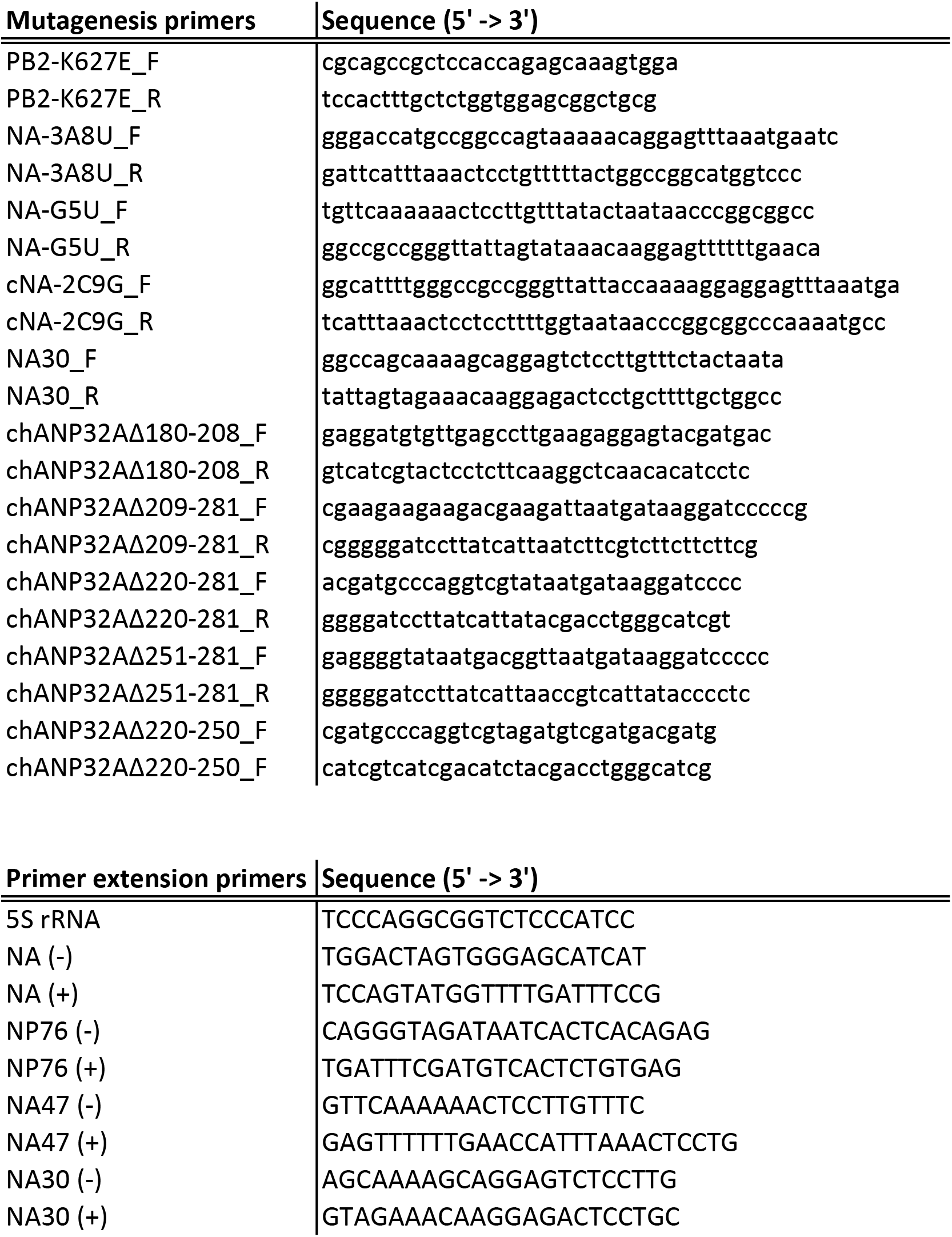
Oligonucleotide sequences.

### Virus infections

Recombinant influenza A/WSN/33 (H1N1) virus was generated using the pHW2000 eight-plasmid reverse genetics system as previously described (51). To generate a recombinant virus with the avian signature lysine-to-glutamic acid point mutation at residue 627 of the PB2 segment, virus rescue was performed with either the wild-type pHW2000-PB2 (WSN) or the mutated pHW2000-PB2-K627E (WSN-K627E) segment.

For viral infections, approximately 1 × 10^6^ human HEK-293T or chicken DF-1 cells were infected with WSN or WSN-K627E at an MOI of 1 in DMEM supplemented with 0.5% FBS.

For analysis of primary viral transcription, infections were performed in the presence of 200 μg/ml cycloheximide at an MOI of 10 in DMEM supplemented with 0.5% FBS.

For viral infections of transiently transfected cells, approximately 1 × 10^6^ human HEK-293T cells were transiently transfected with 1 μg of pCMV-3xFLAG-chANP32A, pCMV-3xFLAG-huANP32A or pCMV-3xFLAG-GFP using Lipofectamine 2000 and Opti-MEM according to the manufacturer’s instructions. 24 h post-transfection, cells were infected at an MOI of 1 in DMEM supplemented with 0.5% FBS.

Total RNA was extracted from all infected samples at 6 h post-infection using TRIzol (Invitrogen) and reconstituted in 20 μl nuclease-free water.

### Viral growth curves

Approximately 1 × 10^6^ HEK-293T cells were transiently transfected with 1 μg of pCMV-3xFLAG-chANP32A or pCMV-3xFLAG-GFP using Lipofectamine 2000 and Opti-MEM according to the manufacturer’s instructions. 24 h post-transfection, cells were infected with influenza A/WSN/33 (H1N1) virus bearing either the wild-type PB2-627K or the avian signature PB2-627E amino acid residue at an MOI of 0.001 in DMEM supplemented with 0.5% FBS. Cell culture media containing viral particles was collected at 6, 24 and 48 h post-infection. The concentration of infectious viral particles (plaque-forming units/ml) was determined by plaque assay on MDCK cells.

### RNP reconstitutions

In order to reconstitute vRNPs in a minigenome assay, approximately 1 × 10^6^ HEK-293T cells were transiently transfected with 1 μg of pcDNA-PA, pcDNA-PB1/pcDNA-PB1a, pcDNA-PB2/pcDNA-PB2-K627E, pcDNA-NP, pCMV-3xFLAG-chANP32A/pCMV-3xFLAG-huANP32A/pCMV-3xFLAG-GFP and a plasmid expressing a vRNA or cRNA segment (pPOLI-NA, pPOLI-cNA, pPOLI-NA-3A8U, pPOLI-G5U, pPOLI-3A8U-G5U, pPOLI-cNA-2C9G) using Lipofectamine 2000 and Opti-MEM according to the manufacturer’s instructions. When using plasmids encoding short truncated vRNA templates (pPOLI-NP76, pPOLI-NA47, pPOLI-NA30), transfections were performed in the absence of pcDNA-NP unless otherwise stated. To ensure that equal amounts of DNA was transfected in all conditions, an empty pcDNA-3a vector was used to balance the transfections. Total RNA was extracted 48 h post-transfection using TRIzol (Invitrogen) and reconstituted in 20 μl of nuclease-free water.

### Primer extension analysis

The accumulation of viral mRNA, cRNA and vRNA was analyzed by primer extension using ^32^P-labelled primers specific for negative- or positive-sense segment 6 RNA as well as 5S rRNA as an internal loading control as described previously (52). Primer sequences are detailed in Table 1. Primer extension products were analyzed by 6% 7 M urea PAGE for full-length RNA templates or 12% 7 M urea PAGE for truncated RNA templates and detected by autoradiography. ImageJ was used to analyze and quantify ^32^P-derived detected signal (53).

### siRNA-mediated knockdown of ANP32A/ANP32B

Approximately 1 × 10^6^ A549 cells were transfected with 20 nM ON-TARGETplus Non-Targeting Control Pool (Dharmacon, D-001810-10-20) or 20 nM siGENOME Human ANP32A siRNA SMARTpool (Dharmacon, M-016060-00-0005) and siGENOME Human ANP32B siRNA SMARTpool (Dharmacon, M-020148-01-0005) using Lipofectamine RNAiMax and Opti-MEM according to the manufacturer’s instructions. 48 h post-transfection, cells were either lysed for Western blot analysis or infected with influenza A virus/WSN/33 (H1N1) virus.

For infections, cells were washed with PBS prior to infection, followed by infection with influenza A/WSN/33 (H1N1) virus at an MOI of 1 in DMEM supplemented with 0.3% BSA for 15 h. Total RNA was extracted using TRIzol (Invitrogen) and reconstituted in 20 μl of nuclease-free water.

### Western blot analysis

For Western blot analysis, cells were lysed in NP-40 lysis buffer containing 1× cOmplete Protease Inhibitor Cocktail (Roche) and 1× Phenylmethylsulfonyl fluoride (Sigma Aldrich) and cleared from the insoluble fraction by centrifugation at 17,000 × *g* for 5 min at 4°C. Samples were analyzed by SDS-PAGE and transferred onto nitrocellulose membranes. Membranes were probed with mouse monoclonal anti-Actin (Thermo Scientific, MS-1295), mouse monoclonal anti-FLAG M2 (Sigma-Aldrich, F1804), rabbit polyclonal anti-PHAP1 (Abcam, ab51013) and rabbit monoclonal anti-PHAPI2 (Abcam, ab200836) antibodies. Primary antibodies were detected using HRP-conjugated secondary anti-mouse (GE Healthcare, NA931V) and anti-rabbit (GE Healthcare, NA934V) antibodies and visualized using Immobilon Western Chemiluminescent HRP substrate kit (Millipore) according to the manufacturer’s instructions.

### RNA-binding assay

Approximately 1 × 10^6^ HEK-293T cells were transiently transfected with 1 μg of pcDNA-PA/pcDNA-PA-TAP, pcDNA-PB1a, pcDNA-PB2/pcDNA-PB2-K627E, pPOLI-NA47 and pCMV-3xFLAG-chANP32A/pCMV-3xFLAG-GFP using Lipofectamine 2000 and Opti-MEM according to the manufacturer’s instructions. 48 h post-transfection, cells were lysed for 1 h at 4°C in 100 μl lysis buffer (50 mM Tris-HCl [pH=8.0], 150 mM NaCl, 0.5% NP-40, 1 mM DTT, 1 mM Phenylmethylsulfonyl fluoride, 1 mM Na_3_VO_4_, 30 mM NaF, 10% glycerol, 1× cOmplete Protease Inhibitor Cocktail). After clarification of lysates from cellular debris by centrifugation (17,000 × *g*, 5 min, 4°C), lysates were incubated for 24 h with equilibrated IgG Sepharose 6 Fast Flow beads at 4°C. Beads were washed three times for 10 min at 4°C with 500 μl wash buffer (10 mM Tris-HCl [pH=8.0], 150 mM NaCl, 0.1% NP-40, 10% glycerol). Beads were collected by centrifugation (17,000 × *g*, 5 min, 4°C) and supernatants were discarded. RNA was extracted from whole cell lysates and IgG Sepharose beads using TRIzol and 20 μg GlycoBlue Coprecipitant (Invitrogen) according to the manufacturer’s instructions. RNA was analysed by primer extension and 12% 7M urea PAGE as described above.

### Recombinant RdRp purification and smFRET

Recombinant influenza A/WSN/33 (H1N1) polymerase subunits containing a PB2 627K, 627E or delta 627 domain and a protein-A tag on the C-terminus of the PA subunit were expressed in HEK-293T cells and purified using IgG-sepharose as described previously (54, 55). RNA and RNA polymerase were prepared and interactions between the molecules measured on a custom-build confocal microscope as described previously. (54, 55)

### cRNA/vRNA stabilization assay

The ability of the viral RdRp to either stabilize nascent cRNA products during primary viral replication or to facilitate replication of both vRNA and cRNA, was tested by transiently transfecting approximately 1 × 10^6^ HEK-293T cells with 1 μg of pcDNA-PA, pcDNA-PB1/pcDNA-PB1a, pcDNA-PB2/pcDNA-PB2-K627E, pcDNA-NP and pCMV-3xFLAG-chANP32A/pCMV-3xFLAG-GFP as indicated using Lipofectamine 2000 and Opti-MEM according to the manufacturer’s instructions. 48 h post-transfection, cells were infected with influenza A/WSN/33 (H1N1) virus in the presence of 5 μg/ml Actinomycin D at an MOI of 5 in DMEM supplemented with 0.5% FBS. Total RNA was extracted 6 h post-infection using TRIzol (Invitrogen) and reconstituted in 20 μl of nuclease-free water. The accumulation of viral mRNA, cRNA and vRNA was analyzed by primer extension using ^32^P-labelled primers specific for negative- or positive-sense segment 6 RNA as well as 5S rRNA as an internal loading control as described previously (52). Primer extension products were analyzed by 6% 7 M urea PAGE in TBE buffer and detected by autoradiography. ImageJ was used to analyze and quantify ^32^P-derived detected signal (53).

## ACKNOWLEDGEMENTS

We thank Ervin Fodor for helpful discussion and advice throughout the duration of this project. This study was supported by Wellcome Trust studentship 102053/Z/13/Z to B.E.N.; National Institutes of Health grants 5R01AI145882 and 5R01AI123155 to B.R.T.; joint Wellcome Trust and Royal Society grant 206579/Z/17/Z to A.t.V., and National Institutes of Health grant R21AI147172 to A.t.V.

